# High-fidelity fast volumetric brain MRI using synergistic wave-controlled aliasing in parallel imaging and a hybrid denoising generative adversarial network

**DOI:** 10.1101/2021.01.07.425779

**Authors:** Ziyu Li, Qiyuan Tian, Chanon Ngamsombat, Samuel Cartmell, John Conklin, Augusto Lio M. Gonçalves Filho, Wei-Ching Lo, Guangzhi Wang, Kui Ying, Kawin Setsompop, Qiuyun Fan, Berkin Bilgic, Stephen Cauley, Susie Y. Huang

## Abstract

**Purpose:** Reducing scan times is important for wider adoption of high-resolution volumetric MRI in research and clinical practice. Emerging fast imaging and deep learning techniques provide promising strategies to accelerate volumetric MRI without compromising image quality. In this study, we aim to leverage an advanced fast imaging technique, wave-controlled aliasing in parallel imaging (Wave-CAIPI), and a novel denoising generative adversarial network (GAN) to achieve accelerated high-fidelity, high-signal-to-noise-ratio (SNR) volumetric MRI.

**Methods:** 3D T_2_-weighted fluid-attenuated inversion recovery (FLAIR) image data were acquired on 33 multiple sclerosis (MS) patients using a prototype Wave-CAIPI sequence (acceleration factor *R*=3×2, 2.75 minutes) and a standard T_2_-SPACE FLAIR sequence (*R*=2, 7.25 minutes). A hybrid denoising GAN entitled “HDnGAN” composed of a 3D generator (i.e., a modified 3D U-Net entitled MU-Net) and a 2D discriminator was proposed to denoise Wave-CAIPI images with the standard FLAIR images as the target. HDnGAN was trained and validated on data from 25 MS patients by minimizing a combined content loss (i.e., mean squared error (MSE)) and adversarial loss with adjustable weight *λ*, and evaluated on data from 8 patients unseen during training. The quality of HDnGAN-denoised images was compared to those from other denoising methods including AONLM, BM4D, MU-Net, and 3D GAN in terms of their similarity to standard FLAIR images, quantified using MSE and VGG perceptual loss. The images from different methods were assessed by two neuroradiologists using a five-point score regarding sharpness, SNR, lesion conspicuity, and overall quality. Finally, the performance of these denoising methods was compared at higher noise levels using simulated data with added Rician noise.

**Results:** HDnGAN effectively denoised noisy Wave-CAIPI images with sharpness and rich textural details, which could be adjusted by controlling *λ*. Quantitatively, HDnGAN (*λ*=10^−3^) achieved low MSE (7.43 ×10^−4^±0.94×10^−4^) and the lowest VGG perceptual loss (1.09×10^−2^±0.18×10^−2^). The reader study showed that HDnGAN (*λ*=10^−3^) significantly improved the SNR of Wave-CAIPI images (4.19±0.39 vs. 2.94±0.24, *P*<0.001), outperformed AONLM (4.25±0.56 vs. 3.75±0.90, *P*=0.015), BM4D (3.31±0.46, *P*<0.001), MU-Net (3.13±0.99, *P*<0.001) and 3D GAN (*λ*=10^−3^) (3.31±0.46, *P*<0.001) regarding image sharpness, and outperformed MU-Net (4.21±0.67 vs. 3.29±1.28, *P*<0.001) and 3D GAN (*λ*=10^−3^) (3.5±0.82, *P*=0.001) regarding lesion conspicuity. The overall quality score of HDnGAN (*λ*=10^−3^) (4.25±0.43) was significantly higher than those from Wave-CAIPI (3.69±0.46, *P*=0.003), BM4D (3.50±0.71, *P*=0.001), MU-Net (3.25±0.75, *P*<0.001), and 3D GAN (*λ*=10^−3^) (3.50±0.50, *P*<0.001), with no significant difference compared to standard FLAIR images (4.38±0.48, *P*=0.333). The advantages of HDnGAN over other methods were more obvious at higher noise levels.

**Conclusion:** HDnGAN provides robust and feasible denoising while preserving rich textural detail in empirical volumetric MRI data and is superior on both quantitative and qualitative evaluation compared to the original Wave-CAIPI images and images denoised using standard methods. HDnGAN concurrently benefits from the improved image synthesis performance of the 3D convolution and the increased number of samples for training the 2D discriminator from a limited number of subjects. Our study supports the use of HDnGAN in combination with modern fast imaging techniques such as Wave-CAIPI to achieve high-fidelity fast volumetric MRI.

## Introduction

High-resolution volumetric brain MRI is widely used in clinical and research applications to provide rich and detailed anatomical information and delineation of structural pathology. For example, 3-dimensional (3D) T_1_-weighted structural MR image volume exhibits high tissue contrast and is therefore routinely used to reconstruct cerebral cortical surfaces for cortical morphometry, analysis and visualization^1–4^. Furthermore, T_2_-weighted fluid-attenuated inversion recovery (FLAIR) imaging is highly sensitive to white matter abnormalities due to its excellent suppression of cerebrospinal fluid signal and is therefore routinely used to characterize lesion pathology in a wide range of neurological disorders^5^. A major barrier to the greater adoption of volumetric MRI in research and clinical protocols is the long acquisition time (typically ~5-7 minutes for 1 mm isotropic resolution and even longer for sub-millimeter resolution^6^), which may lead to subject anxiety and motion artifacts that compromise image quality, especially for children, elderly subjects, and some patient populations who cannot tolerate long scans.

Fast imaging techniques have been increasingly adopted to accelerate volumetric brain MRI. Parallel imaging methods such as sensitivity encoding (SENSE)^7^ and generalized autocalibrating partial parallel acquisition (GRAPPA)^8^ have been adopted by vendors and are used for routine volumetric MRI acquisition. However, these conventional methods can only achieve moderate acceleration factors (i.e., *R*=3-fold along one phase encoding dimension and R=2×2 along both *k_y_* and *k_z_* axes in SENSE and GRAPPA) before suffering from severe image artifacts and noise amplification (i.e., high g-factor). Modern image acquisition and reconstruction methods such as compressed sensing^9^, low-rank modeling of local k-space neighborhoods (LORAKS)^10^, bunched phase encoding (BPE)^11^, 2D controlled aliasing in parallel imaging (CAIPIRINHA)^12^ and wave-controlled aliasing in parallel imaging (Wave-CAIPI)^13^ have been proposed to achieve even higher acceleration factors.

Among these fast imaging methods, Wave-CAIPI is a state-of-the-art technique that employs a corkscrew trajectory with CAIPI shifts in the *k_y_* and *k_z_* directions to efficiently encode k-space^13^. It better utilizes the available degrees of freedom of acquired information in different coils and uniformly spreads the voxel aliasing. Therefore, it can accelerate volumetric MRI by an order of other fast imaging techniques, the signal-to-noise ratio (SNR) 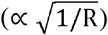 of highly accelerated magnitude with negligible g-factor and image artifact penalties^13–15^. Unfortunately, like images from Wave-CAIPI images is intrinsically lower compared to those acquired without acceleration or with mild acceleration factors due to substantially fewer acquired k-space signals.

Supervised deep learning using convolutional neural networks (CNNs) has superior performance to conventional methods for image restoration tasks such as super-resolution^16^ and denoising^17^ in digital and biomedical imaging. CNNs are remarkably effective in learning the underlying image priors and resolving the complex high-dimensional mapping from low-quality (e.g., low-SNR or low-resolution) images to high-quality images^16–20^, which is an ill-posed inverse problem. Even the simplest CNN for denoising (i.e., DnCNN^17^) achieves superior performance compared to the state-of-the-art block-matching and 3D filtering (BM3D)^21^ denoising method. Recently, it has been shown that DnCNN is also more advantageous in removing noise from sub-millimeter resolution T_1_-weighted structural MR image volumes^22^ compared to the state-of-the-art block matching with 4D filtering (BM4D)^23^ and adaptive optimized nonlocal means (AONLM)^24^. However, a well-known problem of CNNs trained using the voxel-wise error as the loss function is the tendency to generate blurry images that lack realistic textural details in both super-resolution and denoising tasks, even though the mean squared error (MSE), mean absolute error (MAE) or related metrics such as peak SNR (PSNR) can be minimized or maximized^25–27^, respectively.

Generative adversarial networks (GANs)^28^ have been shown to be effective in reducing the blurring effects and recovering realistic textures for digital photography^25,29^, optical coherence tomography^30^, microscopy^31^, x-ray computed tomography^27,32^ and MRI^33–36^. A GAN is composed of two sub-networks, including a generator and a discriminator, which are trained in synchrony to compete against each other. The generator generates high-quality images from the input low-quality images by minimizing the MSE (or the mean of other voxel-wise error measures) between the synthesized and target images, while the discriminator tries to distinguish the synthesized images from the actual target images by maximizing the probability that the synthesized image can be classified as the actual image (i.e., one probability value for one image slice/volume). Essentially, the discriminator serves as a regularizer that forces the generator to synthesize images in the manifold of target images such that they are indistinguishable from target images with similar image features such as image contrast and textural details.

Nonetheless, several barriers limit the performance and feasibility of GANs for denoising empirical volumetric MRI data. Thus far, most studies have demonstrated the efficacy of 2D GANs consisting of a 2D generator and a 2D discriminator on image slices^25,33,37^. The advantage of 2D GANs is that their parameters can be optimized on only a few subjects, because each image volume from a subject provides millions of voxels as training samples for calculating the voxel-wise loss for the generator and hundreds of image slices as training samples for calculating the image-wise loss for the discriminator. However, the image synthesis performance of 2D generators is often limited compared to 3D generators, which can incorporate complementary information from an additional spatial dimension^22,38,39^ (Supplementary Information Figure 1). Moreover, there may be boundary artifacts across synthesized 2D image slices along the cross-sectional direction. Consequently, 3D CNNs are more often adopted for volumetric data^20,22,38–40^ and 3D GANs consisting of a 3D generator and a 3D discriminator have been also proposed^34–36^.

Unfortunately, training a 3D GAN requires data from a large number of subjects since the data of each subject can only provide one image volume as a training sample for the 3D discriminator (or several blocks with smaller size as training samples), which impedes the use of 3D GANs in practice. For example, Chen et al. needed data from 1113 subjects to train a 3D GAN for brain MRI super-resolution^34,36^. Ran et al. had to not only split the whole-brain volumes into small blocks (i.e., 32×32×6 voxels) to increase the training sample number, but also use a sub-optimal network, e.g., a very shallow 3-layer discriminator, which allowed them to train a 3D GAN for MRI denoising with data from 100 subjects^35^. Because of the intensive data requirement, these studies were only performed as a proof of concept on simulated data in which the degraded images were generated by adding noise to high-SNR images or down-sampling high-resolution images from public database^34–36^..

To improve the performance and feasibility of GANs for empirical volumetric MRI data, we propose a hybrid denoising GAN architecture (entitled “HDnGAN”) consisting of a 3D generator and a 2D discriminator which concurrently benefits from the improved image synthesis performance provided by the 3D generator and increased training samples from a limited number of subjects for training the 2D discriminator. Using this new architecture, we demonstrate the efficacy of HDnGAN on empirical Wave-CAIPI T_2_-SPACE FLAIR volumetric data acquired in 33 multiple sclerosis (MS) patients, in distinction to existing studies that only used simulated data^27,34–36^. The systematic and comprehensive comparison and assessment using different quantitative metrics and by two neuroradiologists show that HDnGAN results are similar to standard FLAIR data acquired in longer scan time, and outperform the raw Wave-CAIPI images and those from conventional denoising methods including BM4D and AONLM and a 3D GAN trained on the same training data. In summary, our study increases and proves the superiority and feasibility of GANs in denoising empirical volumetric MRI data and supports its use in combination with modern fast imaging techniques such as Wave-CAIPI adopted in this study to achieve high-fidelity, fast volumetric MRI.

## Methods

### Data acquisition

This study was approved by the Mass General Brigham institutional review board and was HIPAA compliant. With written informed consent, data were acquired in 33 patients (20 for training, 5 for validation, 8 for evaluation) undergoing clinical evaluation for demyelinating disease at the Massachusetts General Hospital as part of a separate clinical validation study of Wave-CAIPI FLAIR compared to standard 3D T_2_-SPACE FLAIR^41^. All patients were scanned on a whole-body 3-Tesla MAGNETOM Prisma MRI scanner (Siemens Healthcare) equipped with a 20-channel head coil.

Standard FLAIR data were acquired using a 3D T_2_-SPACE FLAIR sequence with the following parameters: repetition time = 5000 ms, echo time = 390 ms, inversion time = 1800 ms, flip angle = 120°, 176 sagittal slices, slice thickness = 1 mm, field of view = 256×256 mm^2^, resolution = 1 mm isotropic, bandwidth = 650 Hz/pixel, GRAPPA factor = 2, acquisition time = 7.25 minutes. Wave-CAIPI FLAIR data were acquired using a prototype sequence with matched parameter values except for the following parameters: echo time = 392 ms, bandwidth = 750 Hz/pixel, acceleration factor = 3×2, acquisition time = 2.75 minutes.

### Image processing

Standard FLAIR images were non-linearly co-registered to Wave-CAIPI images using the “*reg_f3d*” function from the NiftyReg software^42,43^, which was initialized with an affine transformation derived from NiftyReg’s “*reg_aladin*” function. The non-linear co-registration slightly adjusted the image alignment locally, which was employed to account for the subtle non-linear shifts of tissue in the images due to factors such as subject bulk motion and the associated changes in distortions due to changing B0 inhomogeneity and gradient nonlinearity distortion during the acquisition of each repetition of data. The resampling used cubic spline interpolation.

Brain masks were created from Wave-CAIPI images using the unified segmentation algorithm in the Statistical Parametric Mapping software^44^. The Wave-CAIPI and standard FLAIR image volumes image volumes were brain masked to exclude regions such as the skull and the background, which are irrelevant to the image content within the brain. To account for subject-to-subject variations in image intensity, the image intensities of the Wave-CAIPI and standard FLAIR image volumes were standardized by subtracting the mean intensity and then dividing by the standard deviation of the image intensities of the voxels within the brain mask in the Wave-CAIPI data.

For each subject, a binary mask was created to exclude parts of the frontal lobe, temporal lobe, cerebellum, and brainstem where the residuals between the Wave-CAIPI and standard FLAIR images were dominated by large image artifacts and geometric distortions rather than noise. These artifactual regions could not be well aligned by image co-registration and would slightly decrease the performance of the CNNs if included since the noise characteristics within these regions are different from those in other brain regions. To create such a mask for each subject, the absolute difference between the standardized Wave-CAIPI and co-registered standard FLAIR images was blurred using a Gaussian kernel with a standard deviation of 2 mm and then binarized using a threshold of 0.04. These parameters were empirically selected and confirmed by visual inspection of all subjects. Only the MSE loss within this mask was used to optimize the generator (i.e., referred to herein as “loss mask”).

### Generative adversarial network

HDnGAN was used to perform image quality transfer from Wave-CAIPI to standard FLAIR images. HDnGAN consisted of a 3D generator and a 2D discriminator. The generator synthesized images similar to standard FLAIR images from the Wave-CAIPI images, while the discriminator tried to distinguish synthesized images from the acquired standard FLAIR images. The generator and the discriminator were trained in synchrony to compete against each other.

A modified 3D U-Net^45^ (denoted as MU-Net) was used as the generator (Fig. 1a) (~2.3 million parameters), which predicted the residuals between the Wave-CAIPI and standard FLAIR image volume (i.e., residual learning). The input and output image volumes consist of *d*×*d*×*d* voxels (*d* = 64 in this study). Specifically, all max pooling, up-sampling, and batch normalization layers were removed, and the number of kernels in each layer was kept constant (*n* = 64). MU-Net created several short paths from early layers to later layers to alleviate the vanishing-gradient problem and strengthen feature propagation with a moderate number of parameters. It represented an intermediate network between a plain network (e.g., VDSR^46^ and DnCNN^17^) without any short paths and a densely connected network (e.g., DenseNet^47^) that comprehensively connects each layer to every other layer.

**Figure 1.**
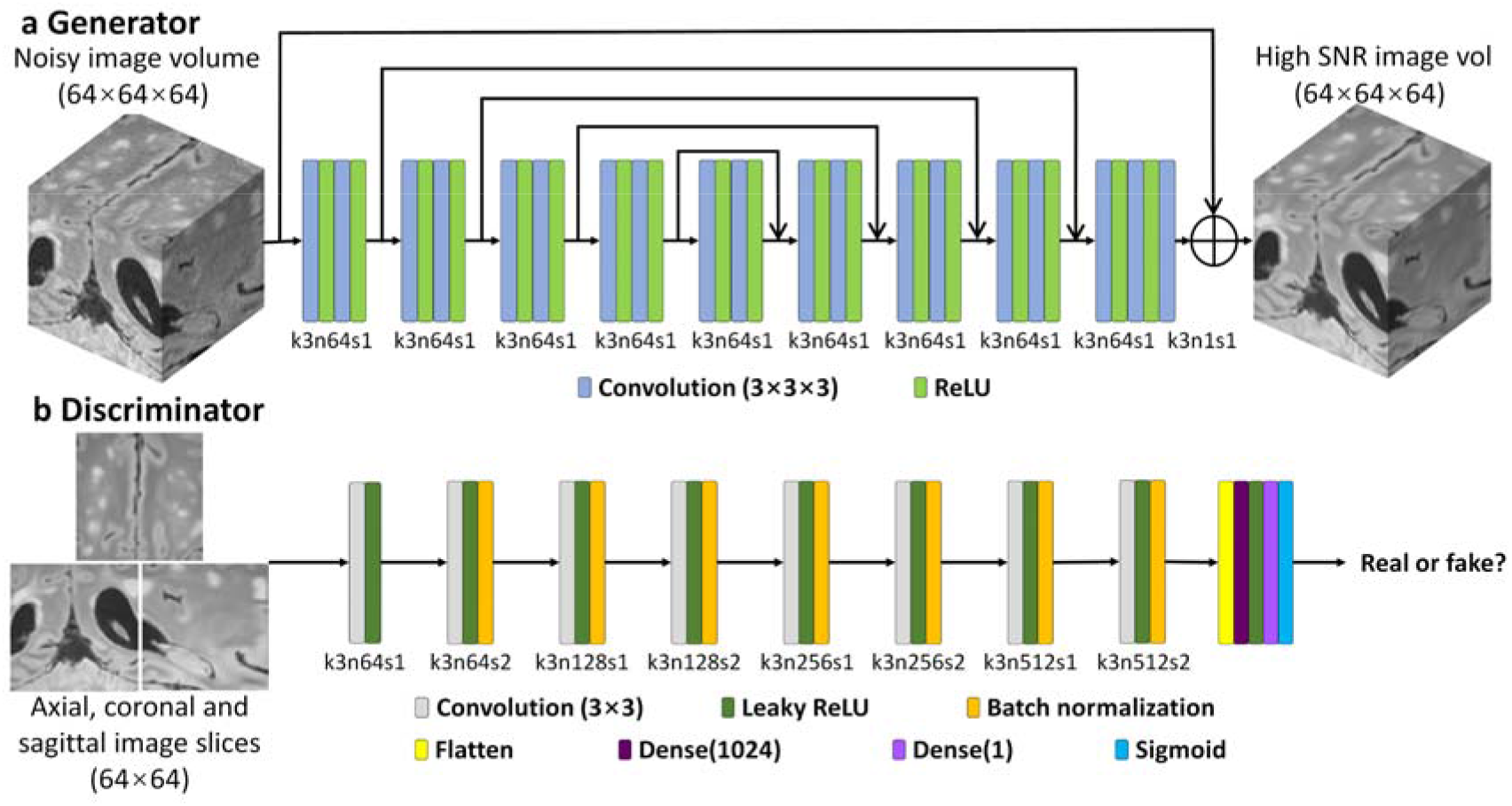
HDnGAN architecture. HDnGAN consists of a 3D generator (a) and a 2D discriminator (b). The 3D generator is modified from 3D U-Net by removing all max-pooling, up-sampling and batch normalization layers and keeping the number of kernels constant (n = 64) across all layers (denoted as MU-Net). The input of the generator is a noisy image volume (64×64×64 voxels). The output of the generator is a synthesized image volume with high signal-to-noise ratio (SNR). The 2D discriminator adopts the discriminator of SRGAN, with spectral normalization incorporated in each layer to stabilize training. The input of the discriminator is an axial, coronal, or sagittal image slice (64×64 pixels) from the image volume synthesized by the generator or the ground-truth high-SNR image volume. The output of the discriminator is a probability value of the input image slice being classified as a real high-SNR image slice. The abbreviation k3n64s1 stands for a kernel size equal to 3×3 for the generator or 3×3×3 for the discriminator, a kernel number equal to 64 and stride equal to 1, and so on.

The discriminator of SRGAN^25^ was adopted (~13.1 million parameters) (Fig. 1b) with spectral normalization incorporated in each layer to stabilize training^48^. The discriminator was designed to classify 2D image slices rather than 3D blocks to increase the number of training samples and ensure that the HDnGAN could be optimized on data from a limited number of subjects. The discriminator used all axial, coronal, and sagittal image slices of the resultant 3D image blocks from the generator as separate samples during the training. In this way, one image volume could provide 3×*d* image slices as training samples to optimize the discriminator parameters, where *d* is the size of the input image volume (*d* = 64 in this study).

### Loss functions

The generator and the discriminator were trained alternately. When the generator was trained, the the generator (θ_G_) such that the synthesized images from the generator were similar to the ground parameters of the discriminator were fixed. The training process tried to optimize the parameters of truth standard FLAIR images. Specifically, the optimizer tried to solve:

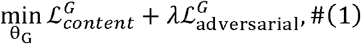

where 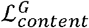 was the voxel-wise MSE between the synthesized images and the standard FLAIR images, 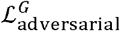 related to the probability that a synthesized image slice could be classified as a real standard FLAIR image, and *λ* determined the contribution of the adversarial loss 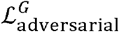 to the total loss 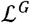, which was a weighted summation of the content loss 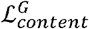 and the adversarial loss 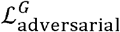. When *λ* = 0, 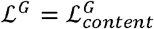, and the HDnGAN was simply the generator (i.e., MU-Net) without the discriminator. When *λ* = ∞, 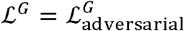. Intermediate *λ*, values achieved mixed content loss and adversarial loss.

The content loss from the generator was defined as the voxel-wise MSE between the synthesized images and the standard FLAIR images, calculated as:

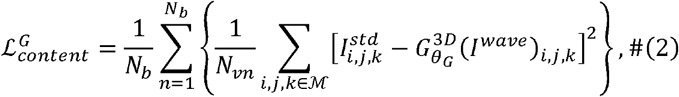

where *I^std^* and *I^wave^* denote standard and Wave-CAIPI image volumes; 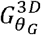 is the 3D generator parametrized by *θ_G_*; *i,j,k* are coordinates within the loss mask 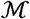 with minimum image artifacts and distortions; *N_vn_* is the number of voxels within the loss mask for each block; *N_b_* is the number of blocks for training.

The adversarial loss was defined as:

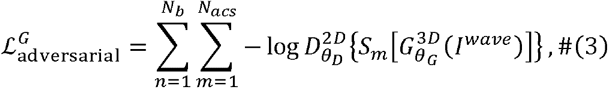

where 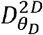 is the 2D discriminator parametrized by *θ_D_* (fixed during the training of the generator); *S_m_*(:) denotes the operation of selecting the *m*th slice out of all *N_acs_* axial, coronal and sagittal slices from the synthesized image block 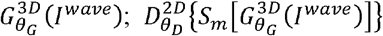 is the probability that an image slice is classified as a real standard FLAIR image.

When the discriminator was trained, the parameters of the generator were fixed. The training process tried to optimize the parameters of the discriminator (θ_D_) such that it could accurately discriminate the When the discriminator was trained, the parameters of the generator were fixed. The training process synthesized and actual images. Specifically, each of the axial, coronal and sagittal slices from the denoised Wave-CAIPI image volumes synthesized by the generator were labeled as 0 and the corresponding real standard FLAIR image slices were labeled as 1. The optimizer tried to solve:

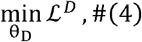

where 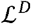 was the binary cross-entropy loss between true labels and predicted labels from the discriminator, defined as:

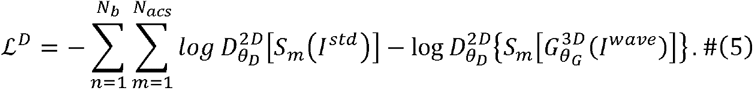

For better gradient behavior, 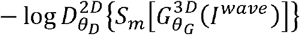 was minimized rather than log {1 − *DθD2DSmGθG3DIwave*^25^.

### HDnGAN training

HDnGAN was implemented using the Keras application program interface (https://keras.io) with a Tensorflow backend (https://www.tensorflow.org) in Python, trained on data from 20 patients, and validated on data from an additional 5 patients using an NVIDIA GeForce RTX 2080 Ti GPU.

Because of the limited GPU memory, the generator was trained on input and output image blocks consisting of 64×64×64 voxels, and the discriminator was trained on input image slices consisting of 64×64 voxels (64×3 axial, coronal, and sagittal image slices from each block). For each subject, 18-27 overlapped blocks were selected, which were evenly distributed within the brain mask with overlap in a sliding window fashion. All training data were flipped along the anatomical left-right direction for augmentation.

Network parameters were optimized as described in equations 1 and 4 using an Adam optimizer with default parameter values except for the learning rates which were set to 5×10^−5^ and 2×10^−4^ for the generator and discriminator respectively. The generator and the discriminator were trained alternately different *λ* values (i.e., 0, 10^−5^, 10^−4^, 10^−3^, 10^−2^, 10^−1^, 1, ∞) were trained and validated for determining the optimal *λ* value, each for 21 epochs when the training of HDnGAN approached convergence (Supplementary Information Fig. 2) and ~24 hours. For *λ* = ∞, the content loss was simply excluded.

### Image denoising using other methods

For comparison, Wave-CAIPI image volumes were also denoised by other state-of-the-art denoising methods, including AONLM and BM4D, and a 3D GAN. The AONLM denoising was performed using the publicly available MATLAB-based software (https://sites.google.com/site/pierrickcoupe/softwares/denoising-for-medical-imaging/mri-denoising/mri-denoising-software) assuming Rician noise with 3×3×3 block and 7×7×7 search volume^24,49^. The BM4D denoising was performed using the publicly available MATLAB-based software (https://www.cs.tut.fi/~foi/GCF-BM3D) assuming Rician noise with the collaborative Wiener filtering and the “modified profile” option with default parameters. The parameters of the 3D GAN were the same as those from HDnGAN, except for that the 3D discriminator classified 64×64×64 generated and ground-truth blocks rather than 64×64 image slices. The 3D GAN with *λ* = 10^−3^ was trained in the same way as the HDnGAN training (e.g., identical training and validation data, 21 epochs for ~24 hours using an Adam optimizer), expect for that the learning rate of the generator was set to 10^−4^. To demonstrate the advantages of 3D generators over 2D generators, a 2D MU-Net for denoising 64×64 image slices was also trained to compare to our 3D MU-Net. The parameters and the training for the 2D MU-Net were identical to those of the 3D MU-Net, except for that the convolutional kernels were 2D.

### Image quality evaluation

Data from 8 separate patients not included in the training were used for evaluation. For each evaluation subject, the optimized HDnGAN and 3D GAN was applied to the extracted image blocks. The resultant denoised image blocks were assembled to create the denoised brain image volume. The overlapping regions of the denoised blocks were averaged.

The group mean and the group standard deviation of the MSE and VGG perceptual loss^25^ across the eight evaluation subjects were used to quantify the similarity between Wave-CAIPI, AONLM-denoised, BM4D-denoised, MU-Net-denoised, 3D GAN-denoised, and HDnGAN-denoised images and standard FLAIR images. For each subject, the MSE was calculated as the mean of the squared differences of the standardized image intensities of all voxels within the brain mask between the denoised and the standard FLAIR image volumes. VGG perceptual loss was calculated as the Euclidean distance between the 4096×1 feature vectors extracted from the denoised and the standard FLAIR image slices using the VGG16 network pre-trained on ImageNet natural image dataset for object classification^50^. For each subject, the VGG loss was calculated for each axial, coronal and sagittal slice from the brain volume and then averaged. The VGG loss closely reflects human perception, and a lower VGG loss value indicates the two images for comparison are visually more similar. For HDnGAN, MSE and VGG perceptual loss were calculated for each weight of adversarial loss.

Images from different methods were also evaluated by two neuroradiologists (C.N., S.C.) based on four image quality metrics, including image sharpness, SNR, lesion conspicuity, and overall quality. For HDnGAN and 3D GAN, results generated using *λ* = 10^−3^ were used for comparison since they had the lowest VGG perceptual loss compared to the standard FLAIR images. The two readers used the standard FLAIR images as the ground truth for evaluation but were blinded to images from the other methods. Images were scored using a five-point scale: 1 nondiagnostic, 2 limited, 3 diagnostic, 4 good, 5 excellent. The group mean and group standard deviation of the scores from each type of images were computed and compared. Moreover, t-tests were performed to assess whether there were significant differences between the image quality scores for different methods. The Benjamini-Hochberg procedure was performed to account for multiple comparisons assuming a 5% false discovery rate.

### Simulation study

A simulation study was performed to demonstrate the efficacy of HDnGAN on noisier images (*I_N_*), which were generated by adding simulated Rician noise to Wave-CAIPI data as:

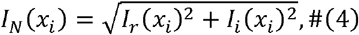

where *I_r_* = *I*_0_(*x_i_*) + *n*_1_(*x_i_*); *I_i_*(*x_i_*) = *n*_2_(*x_i_*); the noise *n* followed a Gaussian distribution with zero mean and standard deviation σ; *I*_0_ was Wave-CAIPI image volumes and *x_i_* represented the coordinates of each voxel. The noise level was described in terms of the mean image intensity 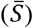 within the brain mask. Noisier data with 2 different noise levels (i.e., 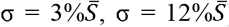) were simulated, which were denoised by AONLM, BM4D, MU-Net, and HDnGAN (*λ* = 10^−3^) as described in previous sections. The denoised results were compared in terms of MSE and VGG perceptual loss.

## Results

The effect of the contribution of adversarial loss on the resultant images is shown in Figure 2. For HDnGAN with low *λ* values (i.e., 0, 10^−5^, 10^−4^) when MSE dominated the loss function, the denoised images were smooth with reduced amount of high-frequency textures (Fig. 2, rows a and b, columns iii-v), as expected. When *λ* gradually increased, more textural details became apparent, and the denoised images became visually more similar to the standard FLAIR images (rows c and d). Images from HDnGAN around *λ* = 10^−3^ (Fig. 2, rows c and d, column i) were visually most similar to the standard FLAIR images.

**Figure 2.**
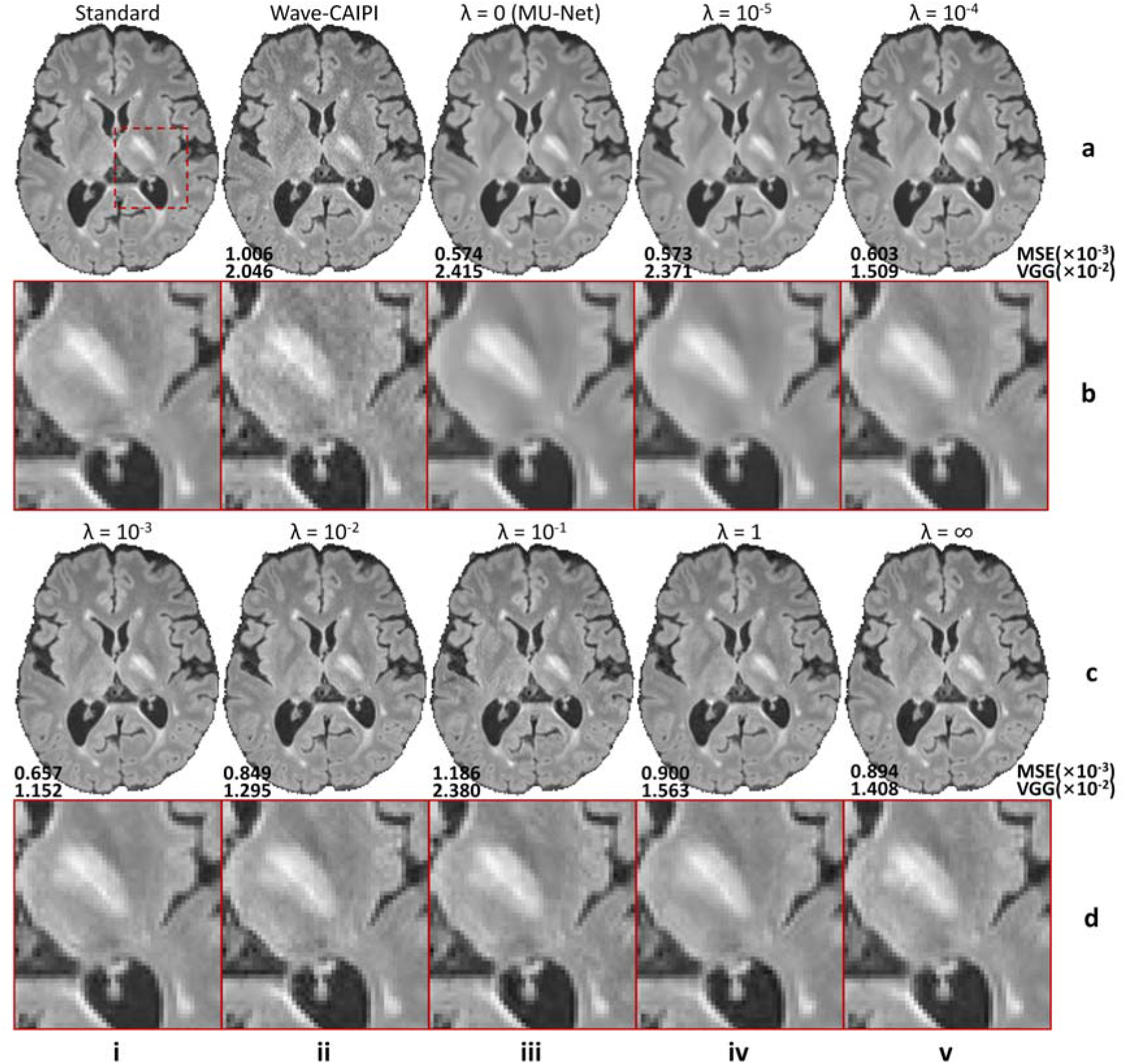
Effects of the adversarial loss on image quality. Representative axial image slices (rows a and c) and enlarged views of the left basal ganglia and left thalamus (rows b and d) from different methods and weights (*λ*) of the adversarial loss in a multiple sclerosis patient for evaluation. For *λ* = 0 (rows a, b, column iii), the training only minimizes the content loss (i.e., voxel-wise mean squared error). In this case, the HDnGAN is effectively the generator (i.e., MU-Net). For *λ* = ∞, the training only minimizes the adversarial loss (rows c, d, column vi). Image similarity metrics including the mean squared error (MSE) and VGG perceptual loss (VGG) are listed to quantify the similarity between images from different methods and the standard FLAIR image.

The optimal *λ* varied accordingly to different image similarity metrics (Fig. 3). In terms of MSE, HDnGAN (*λ* = 0) (i.e., MU-Net) achieved the best performance (5.92×10^−4^ ± 0.56×10^−4^), substantially better than the Wave-CAIPI input images (1.08×10^−3^ ± 0.12×10^−3^) and those generated with adversarial losses (*λ* > 0) (Fig. 3a), which was consistent with the knowledge that denoising CNNs optimized with voxel-wise MSE could minimize MSE but tend to generate blurry images that lack realistic textures (Fig. 2, rows a and b, column iii). When, gradually increased, MSE increased first, reached the maximum at *λ* = 10^−1^ and then decreased. HDnGAN (*λ* = 10^−3^) achieved the best performance in terms of VGG loss (1.09×10^−2^ ± 0.18×10^−2^) (Fig. 3b), improving significantly upon Wave-CAIPI inputs (2.03× 10^−2^ ± 0.33× 10^−2^) and MU-Net (2.45×10^−2^ ± 0.44× 10^−2^), which was consistent with the visual inspection of the denoised images (Fig. 2, rows c and d, column i). As a result, HDnGAN (*λ* = 10^−3^) was used for comparison with other methods. When *λ* gradually increased, VGG perceptual loss first decreased, reached the minimum at *λ* = 10^−3^, then increased until *λ* = 10^−1^, and finally decreased until *λ* = ∞.

**Figure 3.**
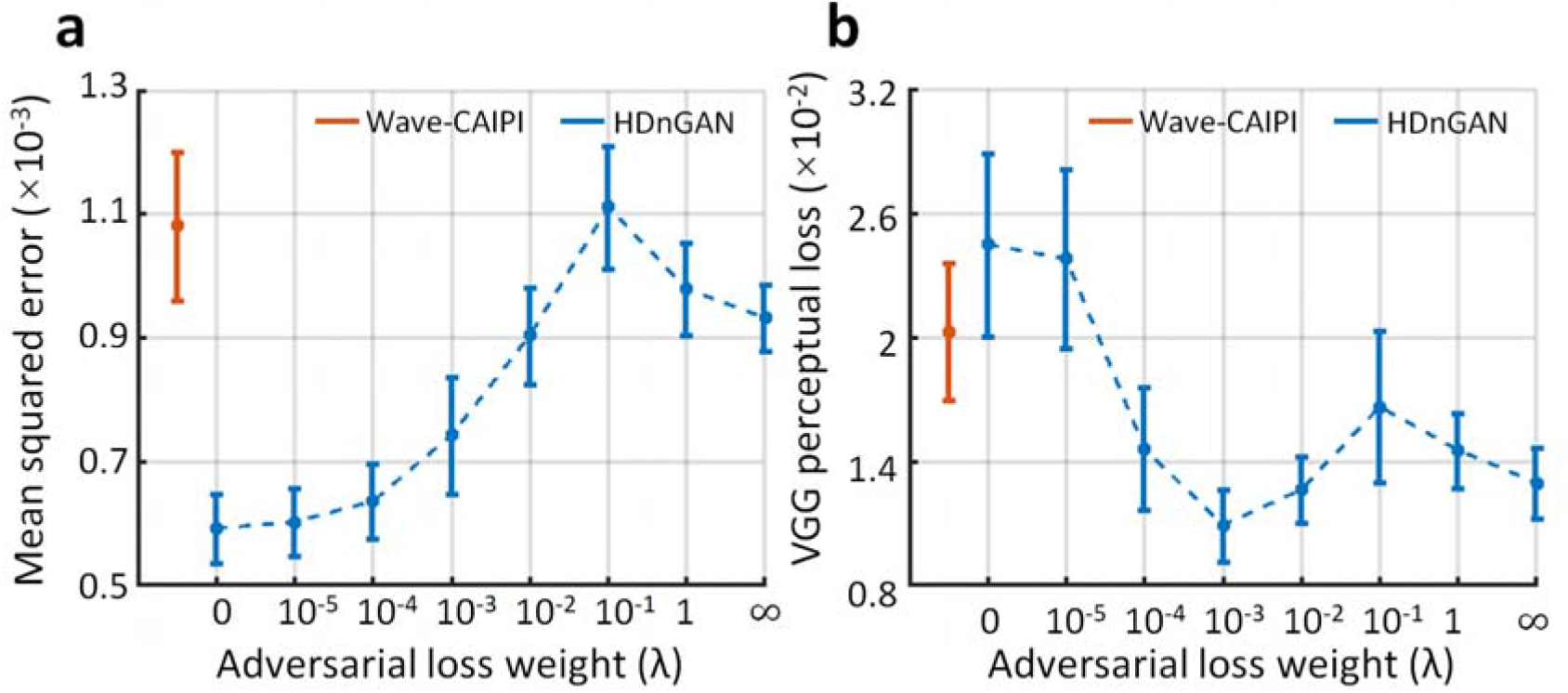
Quantification of the effects of the adversarial loss. The similarity between images derived from different methods and *λ* values and the standard FLAIR images is quantified using the mean squared error (a) and VGG perceptual loss (b). The red and blue dots and error bars represent the group mean and group standard deviation of different metrics for Wave-CAIPI images and results of HDnGAN trained with different weights for the adversarial loss. The metrics were calculated from 8 patients for evaluation.

Denoised images from different methods of a representative subject are shown in Figure 4 for comparison (results from axial and coronal view are available in Supplementary Information Figures 3 and 4). The residual maps between the denoised and standard FLAIR images did not contain anatomical structures (Supplementary Information Fig. 5). All denoised images exhibited higher SNR than the Wave-CAIPI input images. MU-Net-denoised images were the smoothest with reduced amount of texture. The results from the two conventional methods AONLM and BM4D were visually similar and slightly sharper than the MU-Net results. HDnGAN (*λ* = 10^−3^) generated high-quality image volumes with rich and realistic textural details, which were slightly sharper than those from the 3D GAN (*λ* = 10^−3^). This is potentially because the current training and validation data from 25 subjects were not sufficient to optimize the 3D GAN parameters.

**Figure 4.**
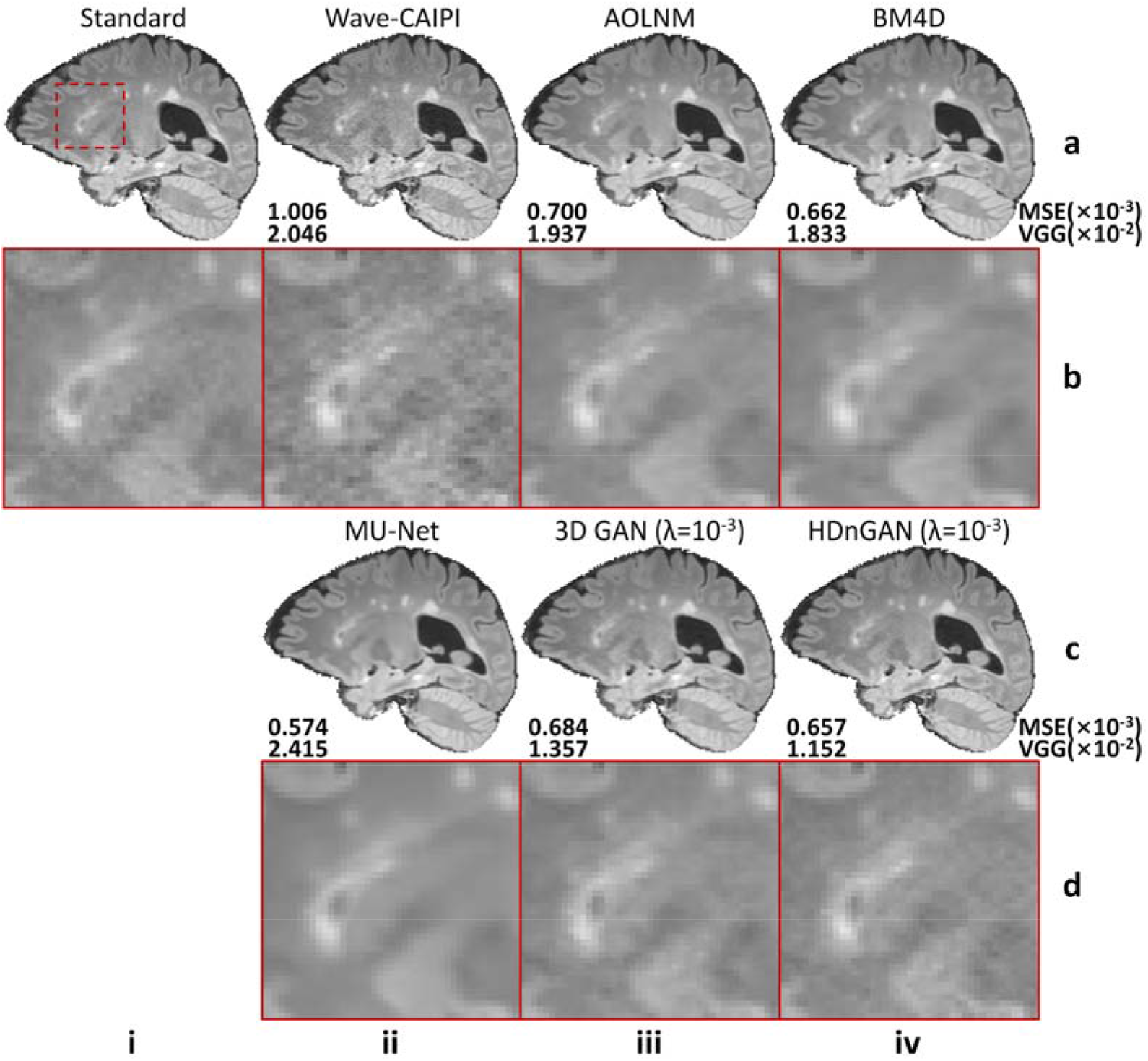
Visual comparison of results from different methods. Representative sagittal image slices (row a and c) and enlarged regions (row b and d) from standard FLAIR data (row a, b, column i), Wave-CAIPI data (row a, b, column ii), AONLM-denoised results (row a, b, column iii), BM4D-denoised results (row a, b, column iv), MU-Net results (row c, d, column ii), 3D GAN (*λ* = 10^−3^) results (row c, d, column iii) and HDnGAN (*λ* = 10^−3^) results (row c, d, column iv) from one evaluation subject. Image similarity metrics including the mean squared error (MSE) and VGG perceptual loss (VGG) are listed to quantify the similarity between images from different methods and the standard FLAIR image.

The group mean and the group standard deviation of the MSE and VGG loss across the eight 8 evaluation subjects were listed in Table 1. Among all methods for comparison, MU-Net-denoised images achieved the best MSE (5.92×10^−4^ ± 0.56×10^−4^) but also the worst VGG loss (2.45×10^−2^ ± 0.44×10^−2^). HDnGAN (*λ* = 10^−3^) generated images with low MSE (7.43×10^−4^ ± 0.94×10^−4^), which was comparable to those from BM4D (7.18×10^−4^ ± 0.75×10^−4^) and AONLM (7.49×10^−4^ ± 0.76×10^−4^) and lower than that from 3D GAN (*λ* = 10^−3^) (7.75×10^−4^ ± 0.95×10^−4^). HDnGAN (*λ* = 10^−3^) achieved the best VGG loss (1.09×10^−2^ ± 0.18×10^−2^), which was substantially lower than those from BM4D (1.72×10^−2^ ± 0.23×10^−2^), AONLM (1.69×10^−2^ ± 0.27×10^−2^) and 3D GAN (*λ* = 10^−3^) (1.35×10^−2^ ± 0.18×10^−2^). Finally, HDnGAN (*λ* = 10^−3^) outperformed 3D GAN (*λ* = 10^−3^) in terms of both MSE (7.43×10^−4^ ± 0.94×10^−4^ vs. 7.75×10^−4^ ± 0.95×10^−4^) and VGG loss (1.09×10^−2^ ± 0.18×10^−2^ vs. 1.35×10^−2^ ± 0.18×10^−2^).

**Table 1.** Quantitative comparison of results from different methods. The group mean and group standard deviation of image similarity metrics (mean ± std) including the mean squared error and VGG perceptual loss are listed to quantify the similarity between images from different methods and the standard FLAIR image. The methods with the best performance for each metrics are marked in bold. The metrics were calculated from eight patients for evaluation.

| Method | Mean squared error ($\times 10^{-3}$) | VGG perceptual loss ($\times 10^{-2}$) |
| --- | --- | --- |
| Wave-CAIPI | 1.082±0.120 | 2.031±0.334 |
| AONLM | 0.749±0.076 | 1.687±0.271 |
| BM4D | 0.718±0.075 | 1.716±0.230 |
| MU-Net | **0.592±0.056** | 2.453±0.444 |
| 3D GAN ($\lambda=10^{-3}$) | 0.775±0.095 | 1.349±0.179 |
| HDnGAN ($\lambda=10^{-3}$) | 0.743±0.094 | **1.091±0.175** |

The results of the reader study (Fig. 5, Supplementary Information Table 1) demonstrated that HDnGAN (*λ* = 10^−3^) significantly outperformed AONLM (4.25 ± 0.56 vs. 3.75 ± 0.90, *P* = 0.015), BM4D (4.25 ± 0.56 vs. 3.31 ± 0.46, *P* < 0.001), MU-Net (4.25 ± 0.56 vs. 3.13 ± 0.99, *P* < 0.001) and 3D GAN (*λ* = 10^−3^) (4.25 ± 0.56 vs. 3.31 ± 0.46, *P* < 0.001) in terms of image sharpness. HDnGAN (*λ* = 10^−3^) also significantly improved the input Wave-CAIPI images’ SNR (4.19 ± 0.39 vs. 2.94 ± 0.24, *P* < 0.001) and outperformed 3D GAN (*λ* = 10^−3^) (4.19 ± 0.39 vs. 3.75 ± 0.43, *P* = 0.004). The lesion conspicuity of the generated images from HDnGAN (*λ* = 10^−3^) was significantly better than those from MU-Net (4.21 ± 0.67 vs. 3.29 ± 1.28, *P* < 0.001) and 3D GAN (*λ* = 10^−3^) (4.21 ± 0.67 vs. 3.5 ± 0.82, *P* = 0.001). The overall scores from HDnGAN (*λ* = 10^−3^) were significantly higher than those from Wave-CAIPI (4.25 ± 0.43 vs. 3.69 ± 0.46, *P* = 0.003), BM4D (4.25 ± 0.43 vs. 3.50 ± 0.71, *P* = 0.001), MU-Net (4.25 ± 0.43 vs. 3.25 ± 0.75, *P* < 0.001) and 3D GAN (*λ* = 10^−3^) (4.25 ± 0.43 vs. 3.50 ± 0.50, *P* < 0.001), with no significant difference compared to the standard FLAIR images (4.25 ± 0.43 vs. 4.38 ± 0.48, *P* = 0.333).

**Figure 5.**
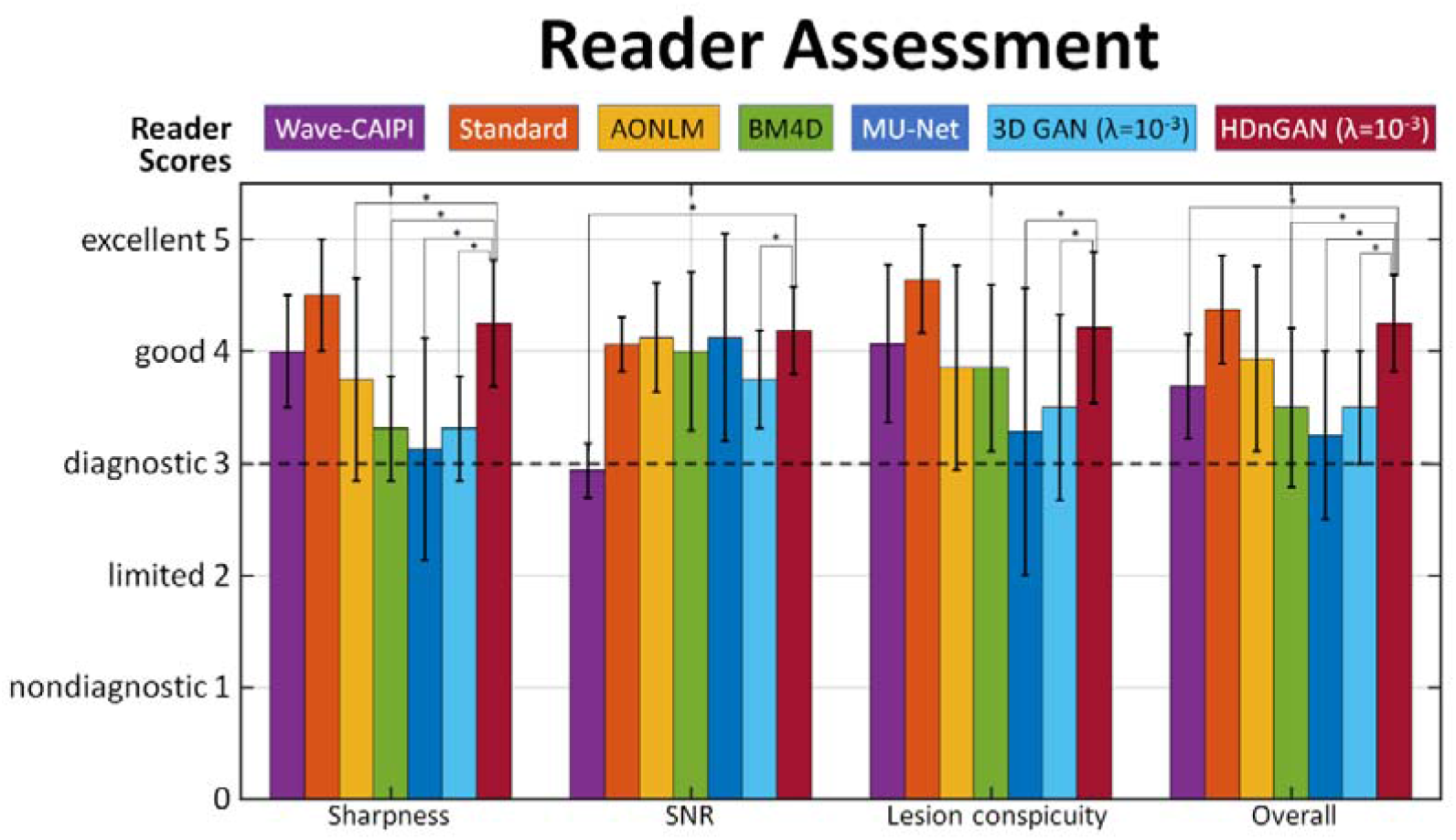
Multi-reader assessment of results from different methods. The group mean and group standard deviation of image quality (sharpness, SNR, lesion conspicuity, and overall quality) scores (1 nondiagnostic, 2 limited, 3 diagnostic, 4 good, 5 excellent) from two radiologists for the standard, Wave-CAIPI, AONLM, BM4D, MU-Net, 3D GAN (*λ* = 10^−3^), and HDnGAN (*λ* = 10^−3^) images of eight evaluation subjects for evaluation. Comparisons between scores of HDnGAN (*λ* = 10^−3^) and scores of other methods are denoted surviving multiple comparisons correction at an FDR threshold of 0.05 are denoted with an asterisk.

The advantages of HDnGAN became more obvious for noisier data in the simulation study (Fig. 6, Table 2). With moderate noise 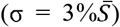 added, the blurring effects of AONLM, BM4D, and MU-Net became more obvious (Fig. 6, rows a and b, columns iii-v), while HDnGAN (*λ* = 10^−3^) improved the image SNR and preserved realistic textures (Fig. 6, rows a and b, columns vi), achieving the lowest VGG perceptual loss (1.336×10^−2^ ± 0.186×10^−2^). MU-Net achieved the lowest MSE (0.703 ×10^−3^ ± 0.065×10^−3^). At a very high noise level 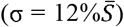, AONLM-denoised and BM4D-denoised images exhibited obvious errors in anatomical structures, e.g., with obscured contrast between the gray matter and white matter (Fig. 6, rows c and d, columns iii, iv). MU-Net still achieved the lowest MSE (1.324×10^−3^ ± 0.039×10^−3^) but the output images were very blurry (Fig. 6, rows c and d, columns v). HDnGAN (*λ* = 10^−3^) removed noise and recovered most signals with acceptable visual quality (Fig. 6, rows c and d, columns vi) and achieved the best VGG perceptual loss (2.834×10^−2^ ± 0.425×10^−2^).

**Figure 6.**
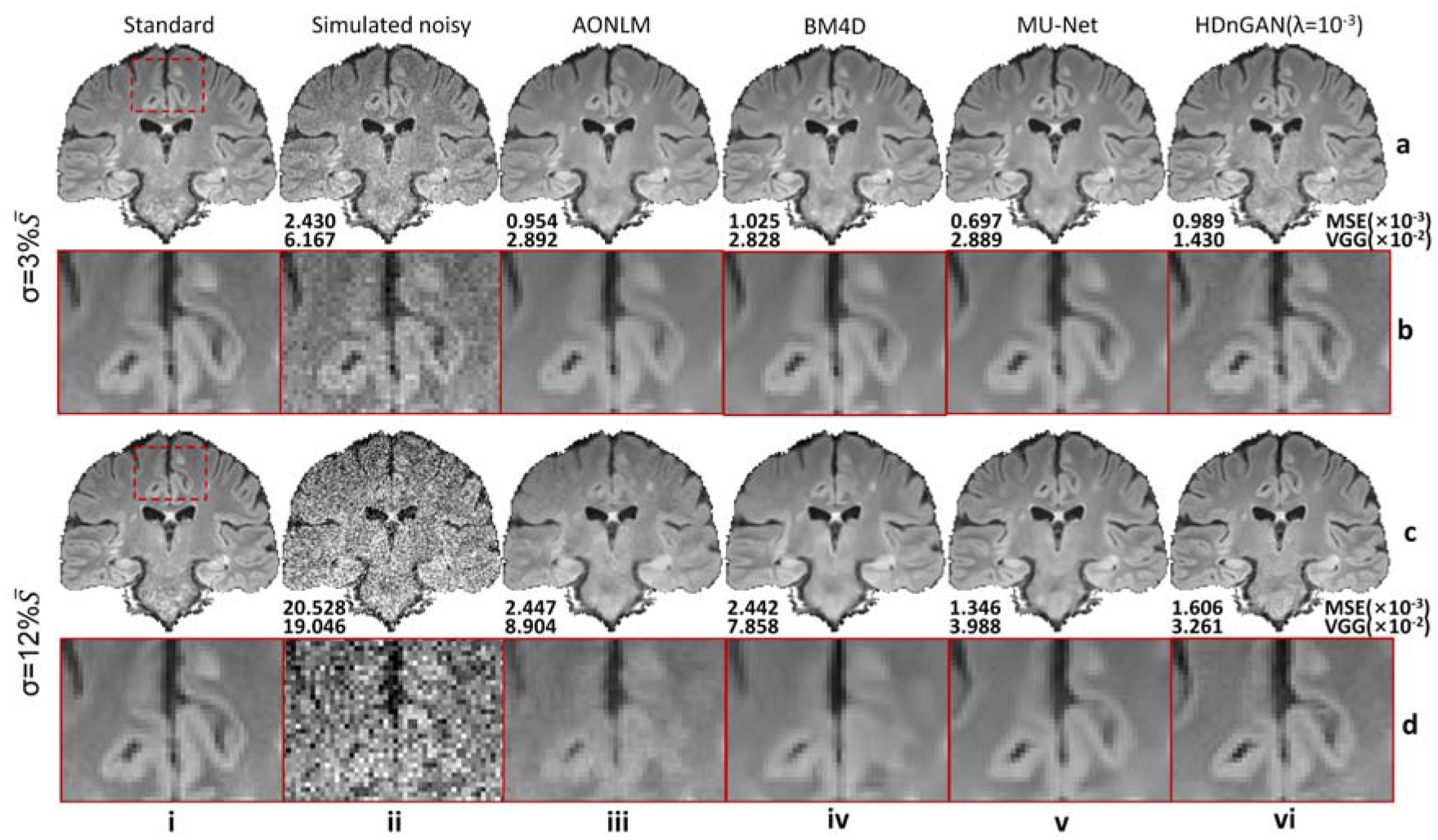
Visual comparison of results from different methods at higher noise levels. Representative coronal image slices (row a and c) and enlarged regions (row b and d) from the standard FLAIR data (column i), simulated noisy data (column ii), AONLM-denoised data (column iii), BM4D-denoised data (column iv), MU-Net-denoised data (column v) and HDnGAN-denoised (*λ* = 10^−3^) data (column v) for different levels of simulated noise. The noise level is described in terms of the standard deviation (σ) of the Gaussian noise components of the Rician noise added to Wave-CAIPI data. 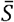 denotes the mean image intensity within the brain mask. Metrics including the mean squared error (MSE) and VGG perceptual loss (VGG) are listed to quantify the similarity between images from different methods and the standard FLAIR images of the patient.

**Table 2.** Quantitative comparison of results from different denoising methods at higher noise levels. The group mean and group standard deviation of image similarity metrics (mean ± std) including the mean squared error and VGG perceptual loss are listed to quantify the similarity between images from different methods and the standard FLAIR image at higher noise levels. The noise level is described in terms of the standard deviation (σ) of the Gaussian noise components of the Rician noise added to Wave-CAIPI data. 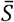 denotes the mean image intensity within the brain mask. The methods with the best performance for each metrics of each noise level are marked in bold. The metrics were calculated from eight patients for evaluation.

| Noise level | Method | Mean squared error ($\times 10^{-3}$) | VGG perceptual loss ($\times 10^{-2}$) |
| --- | --- | --- | --- |
| $\sigma = 3\% \bar{S}$ | Simulated noisy | 2.306±0.124 | 5.701±0.391 |
| | AONLM | 0.940±0.072 | 2.700±0.399 |
| | BM4D | 0.960±0.072 | 2.779±0.388 |
| | MU-Net | **0.703±0.065** | 2.785±0.483 |
| | HDnGAN ($\lambda = 10^{-3}$) | 0.917±0.050 | **1.336±0.186** |
| $\sigma = 12\% \bar{S}$ | Simulated noisy | 20.544±0.111 | 18.523±1.071 |
| | AONLM | 2.228±0.144 | 8.031±0.510 |
| | BM4D | 2.126±0.183 | 6.495±0.913 |
| | MU-Net | **1.324±0.039** | 3.637±0.486 |
| | HDnGAN ($\lambda = 10^{-3}$) | 1.550±0.043 | **2.834±0.425** |

## Discussion

This study leveraged data acquired using the state-of-the-art fast imaging method Wave-CAIPI and machine learning technology GAN to achieve fast volumetric MRI with high-fidelity image quality similar to the standard images acquired in 2.6-times longer scan time. The proposed HDnGAN not only improves SNR but also recovers realistic textures, the richness of which can be controlled by adjusting adversarial loss contributions. Quantitatively, HDnGAN (*λ* = 0), i.e., MU-Net, achieves the lowest MSE while HDnGAN (*λ* = 10^−3^) achieves the lowest VGG perceptual loss compared to the standard FLAIR images among all denoising methods, including AONLM, BM4D and the 3D GAN trained on limited training data. Reader assessment demonstrates radiologists’ preference for images from HDnGAN (*λ* = 10^−3^) over raw Wave-CAIPI images and denoised images from AONLM, BM4D, MU-Net, and the 3D GAN (*λ* = 10^−3^).

Image denoising provides a complimentary approach for improving the intrinsically low SNR of modern fasting MRI techniques. Even though the state-of-the-art fast imaging methods such as the Wave-CAIPI adopted in our study can achieve high acceleration rates with negligible noise amplification and structural artifacts penalties, the SNR of highly accelerated images suffers from the inherent 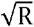 penalty, which substantially benefits from image denoising using frameworks such as compressed sensing^9,33^ and LORAKS^10^ as well as CNNs. In our study, the denoising step by CNNs is cast as a stand-alone post-processing step with several advantages. First, it only takes in reconstructed images from the MRI scanner as inputs, without any need to intervene in the current imaging flow or handle the very large k-space data from individual coils. Therefore, our proposed method can be easily incorporated into existing software packages for visualization and analysis, such as real-time visualization of resultant image on the console computers, for immediate benefits. Second, the denoising process is independent of the imaging process, such that HDnGAN can be applied to noisy images from any MRI contrast, sequences or reconstruction methods (e.g., those provided by the vendor, diffusion-weighted imaging or susceptibility weighted imaging) and imaging modalities (e.g., x-ray computed tomography, positron emission tomography). Third, denoising images also facilitates the acquisition of the training data as reconstructed images from a wider range of MRI scanners (e.g., those in hospitals) or the use of legacy image data as the training data, which is especially critical for GANs that require much more training data than normal CNNs. This also greatly facilitates clinicians and neuroscientists who most often use reconstructed images from MRI scanners.

Thus far, only 2D GANs (consisting of a 2D generator and a 2D discriminator) or 3D GANs (consisting of a 3D generator and a 3D discriminator) have been proposed, which have their own pros and cons. 2D GANs can be trained on data from a few subjects, since each data volume provides millions of voxels and hundreds of image slices as training samples for the generator and the discriminator, respectively. However, the image synthesis performance of 2D generators is limited compared to 3D generators, which is clearly demonstrated using a 2D MU-Net (trained on sagittal image slices) and a 3D MU-Net in Supplementary Information Figure 1. In the images from the 2D MU-Net, the lesion contrast is reduced in the sagittal plane and some subtle lesions become undetectable (Supplementary Information Fig. 1 red arrows) and the transition between 2D images slices is not smooth in the axial (Supplementary Information Fig. 1 yellow arrows) and coronal view (Supplementary Information Fig. 1 blue arrows) such that the lesion geometry is obscured. Quantitatively, the 2D MU-Net exhibits higher MSE compared to the 3D MU-Net (6.64×10^−4^ ± 0.67×10^−4^ vs. 5.92×10^−4^ ± 0.56×10^−4^ for the eight evaluation patients).

On the other hand, the 3D discriminator in a 3D GAN requires substantially more data for training because its loss is computed for a single probability value that classifies the input image volume and the data from a subject can only provide one image volume or several image blocks (e.g., 18-27 blocks of 64×64×64 voxels). In theory, 3D GANs are superior, if trained and validated on sufficient data, since the 3D discriminator has more information to distinguish the 3D synthesized and real image blocks. In previous simulation studies which used 1113 subjects or 100 subjects for training and validation, 3D GANs are indeed demonstrated effective for image super-resolution and denoising^34–36^. However, our study shows that the 3D GAN cannot compare to the proposed HDnGAN (Figs. 4, 5, Table 1) if trained and validated on the data from only 25 MS patients. Presumably this is because the 10-layer 3D discriminator (13.1 million parameters) cannot be optimized using only ~650 blocks from 25 MS patients. Acquiring training data on much more subjects, e.g., 100+ MS patients, would be challenging in practice. This is an important reason that previous studies have only proven the principle of 3D GANs using simulation data but have never demonstrated their efficacy on empirical data.

The hybrid architecture of HDnGAN with a 3D generator and a 2D discriminator facilitates HDnGAN to be well-trained on limited empirical data (e.g., 25 patients in our study), which makes it possible to use GANs in practice. The use of 2D discriminator in HDnGAN substantially increases the training samples by 64×3 times by classifying all axial, coronal, and sagittal slices (i.e., ~650 blocks × 64 × 3 image slices), which provides a practical solution for concurrent benefits from a 3D generator and a 2D discriminator and also mimics the visual inspection by radiologists who read 2D images from different views. The results confirm HDnGAN’s superiority to state-of-the-art 3D GAN in objective evaluation including data consistency (i.e., MSE) and visual quality (i.e., VGG), and subjective evaluation including sharpness, SNR, lesion conspicuity, and overall quality as it alleviates the overfitting problem for the 3D discriminator when training data are limited.

We also systematically characterized the effects of adversarial loss on resultant image sharpness. Overall, larger adversarial loss weight leads to images with more textural details (Fig. 2) with lower VGG perceptual loss but higher MSE (Fig. 3). In order to demonstrate this effect, we did not use the VGG perceptual loss in the optimization as in many previous studies^25,27,35,37^, which confounds the origin of the resultant textural details. Clarifying this effect not only facilitates the selection of optimal weight of the adversarial loss (i.e., *λ*), but also enables an elegant way to control the output image sharpness, which could address the needs of radiologists who may have different preferences for textures’ richness. In our study, we select *λ* = 10^−3^ which generates output images that are most similar to the standard FLAIR images as quantified by the VGG perceptual loss (even though VGG was trained on natural images from ImageNet database) and by visual inspection of radiologists (Fig. 2). In practice, *λ* could be selected or adjusted by radiologists according to their preferences.

Previous studies often perform a linear blending of the GAN generator network parameters^29^ or output images of models optimized using only the MSE loss and the adversarial loss^29,36^ to achieve different sharpness levels for image super-resolution. However, these two methods are sub-optimal as shown in Supplementary Information Figures 6 and 7. Visually, the network parameter blending approach fails to denoise Wave-CAIPI images effectively, while the textures from the image blending method are less realistic than those from HDnGAN (*λ* = 10^−3^) (Supplementary Information Fig. 6). The VGG perceptual losses of both methods are higher than HDnGAN (*λ* = 10^−3^) results (Supplementary Information Fig. 7), demonstrating the superiority of our method on controlling resultant image sharpness.

The results of reader study confirm the superiority of HDnGAN as HDnGAN (*λ* = 10^−3^) achieves the highest mean score among all denoising methods for comparison. The mean overall score of HDnGAN (*λ* = 10^−3^) is significantly higher than those of Wave-CAIPI, BM4D, MU-Net, and 3D GAN (*λ* = 10^−3^), with no significant difference compared to standard FLAIR images. The mean overall score from HDnGAN (*λ* = 10^−3^) is also higher than that of AONLM but the comparison is not significant. The large standard deviation of overall scores for images from AONLM contribute to the insignificance, which indicates the two radiologists have varied opinions on the resultant images from AONLM.

HDnGAN and the approach to control the output image sharpness can be simply extended for different noise levels, MRI sequences and contrasts (e.g., diffusion-weighted MRI), imaging tasks (e.g., super-resolution), and CNN-based image reconstruction from k-space data, and different biomedical imaging modalities. The acceleration factor of the Wave-CAIPI data used for our experiment was chosen not very high (3×2) due to the use of the 20-channel head coil. For 32-channel or 64-channel^51^ coils, HDnGAN could be employed to denoise Wave-CAIPI images with 10× and even higher acceleration to further reduce the scan time to within a minute. As our simulation experiments suggest, the advantages of HDnGAN are more obvious at a higher noise level (Fig. 6, Table 2). Further, HDnGAN can be used for a variety of other Wave-CAIPI-accelerated 3D sequences beyond T_2_-weighted FLAIR^41^, including 3D T_1_-weighted magnetization prepared rapid gradient echo (MPRAGE)^15,52^, susceptibility-weighted imaging^53^, and T_1_-weighted SPACE^54,55^, as well as images reconstructed using other accelerated methods such as compressed sensing and LORAKS^10,56^. Finally, the generator of HDnGAN can be replaced with any CNN (e.g., variational network^57^) that reconstructs images directly from k-space data in order to recover image sharpness and realistic textural details in the reconstructed images.

## Conclusion

This study proposes a synergistic Wave-CAIPI and GAN method for high-fidelity fast FLAIR imaging. The proposed hybrid denoising GAN, entitled HDnGAN, improves the SNR of highly accelerated Wave-CAIPI images while preserving realistic textural details. HDnGAN benefits from improved image synthesis performance from the 3D generator and increased training samples for training the 2D discriminator on empirical data from 25 MS patients. HDnGAN (*λ* = 10^−3^) generates images most similar to high-quality images acquired in longer scan times, with the lowest VGG perceptual loss and higher preference by neuroradiologists than Wave-CAIPI images and those denoised by AONLM, BM4D, MU-Net, and 3D GAN (*λ* = 10^−3^).

## Supplementary Information

**Supplementary Information Figure 1.**
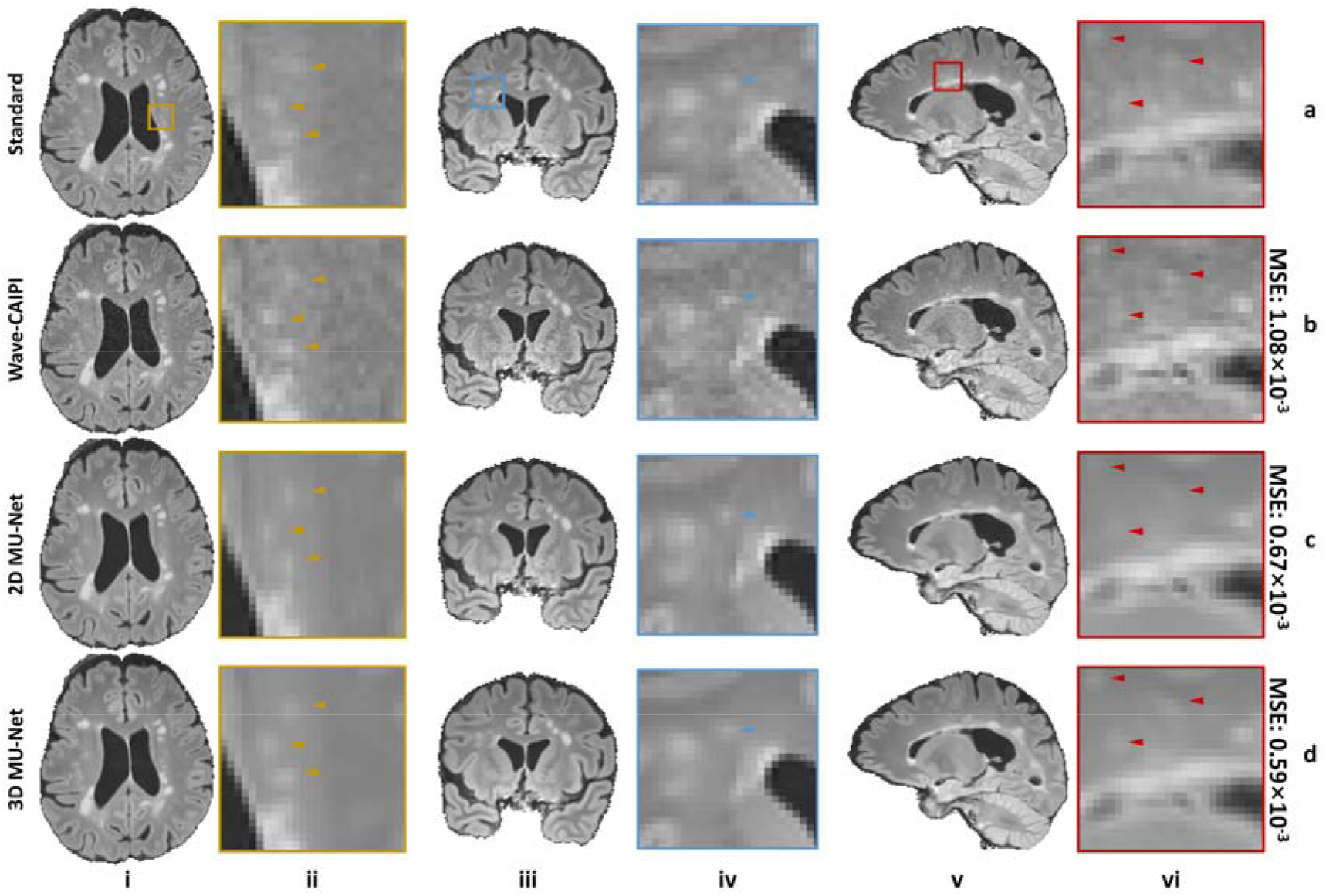
Comparison of 2D and 3D generator. Representative axial (column i), coronal (column iii), and sagittal (column v) image slices and enlarged regions (columns ii, iv and vi) from standard FLAIR data (row a), Wave-CAIPI data (row b), 2D MU-Net-denoised results (row c) and 3D MU-Net-denoised results (row d) from one evaluation subject. The mean squared error (MSE) for the subject is listed to quantify the similarity between images from different methods and the standard FLAIR image.

**Supplementary Information Figure 2.**
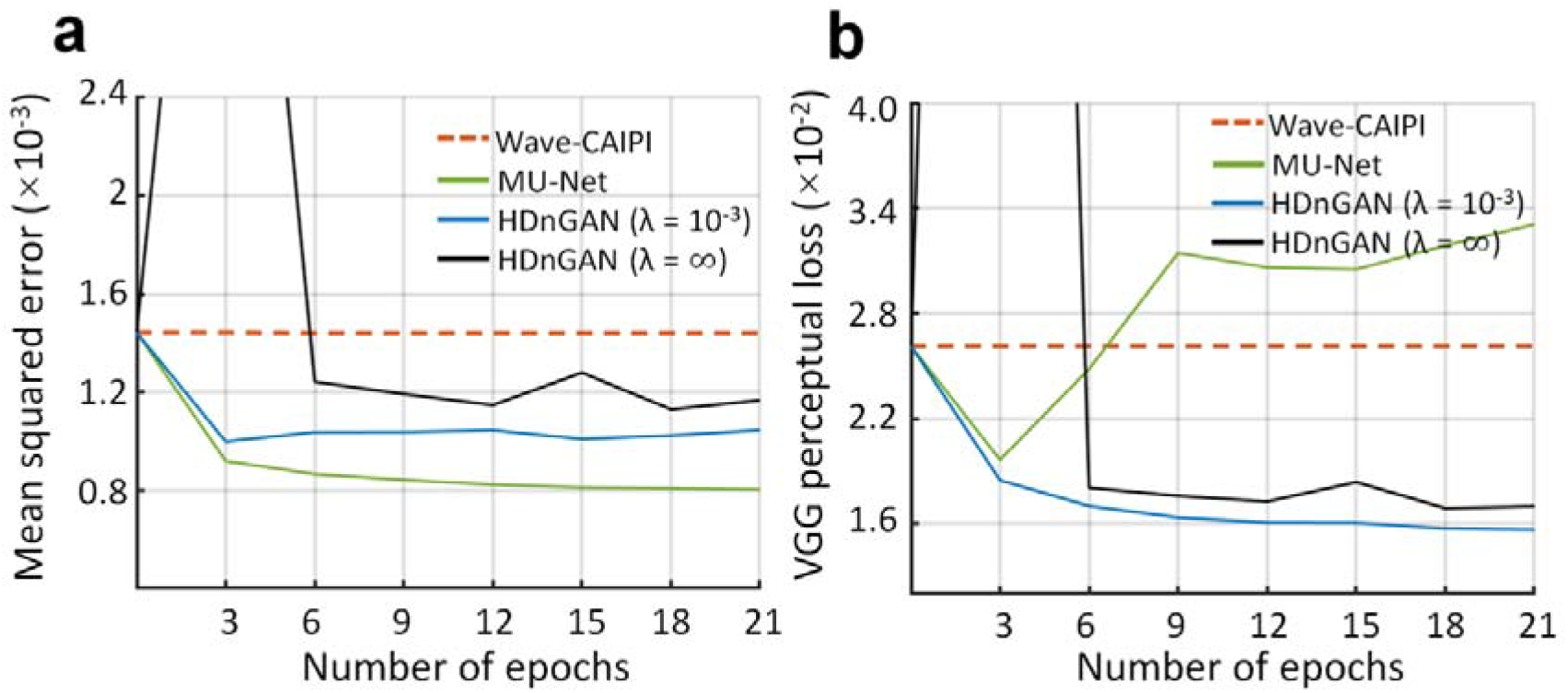
Training convergence. The similarity between standard FLAIR images and Wave-CAIPI images (red dashed lines) as well as resultant images from HDnGAN trained with different weights for the adversarial loss (green, blue and black lines) at different epochs during the training is quantified using the mean squared error (a) and VGG perceptual loss (b). The metrics were calculated from five patients for validation.

**Supplementary Information Figure 3.**
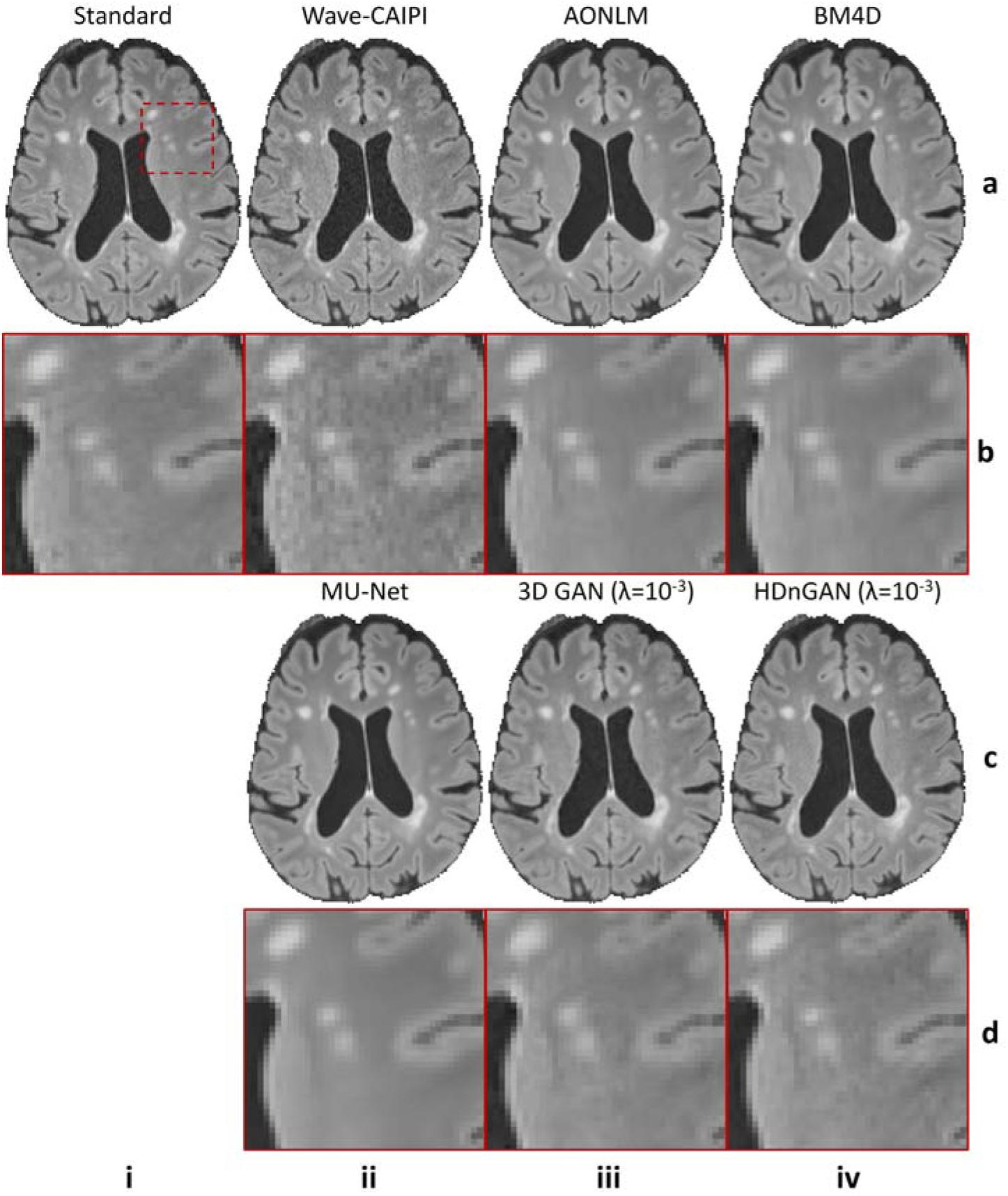
Visual comparison of axial image slices from different methods. Representative axial image slices (row a and c) and enlarged regions (row b and d) from standard FLAIR data (row a, b, column i), Wave-CAIPI data (row a, b, column ii), AONLM-denoised results (row a, b, column iii), BM4D-denoised results (row a, b, column iv), MU-Net results (row c, d, column ii), 3D GAN (*λ* = 10^−3^) results (row c, d, column iii) and HDnGAN (*λ* = 10^−3^) results (row c, d, column iv) from one evaluation subject.

**Supplementary Information Figure 4.**
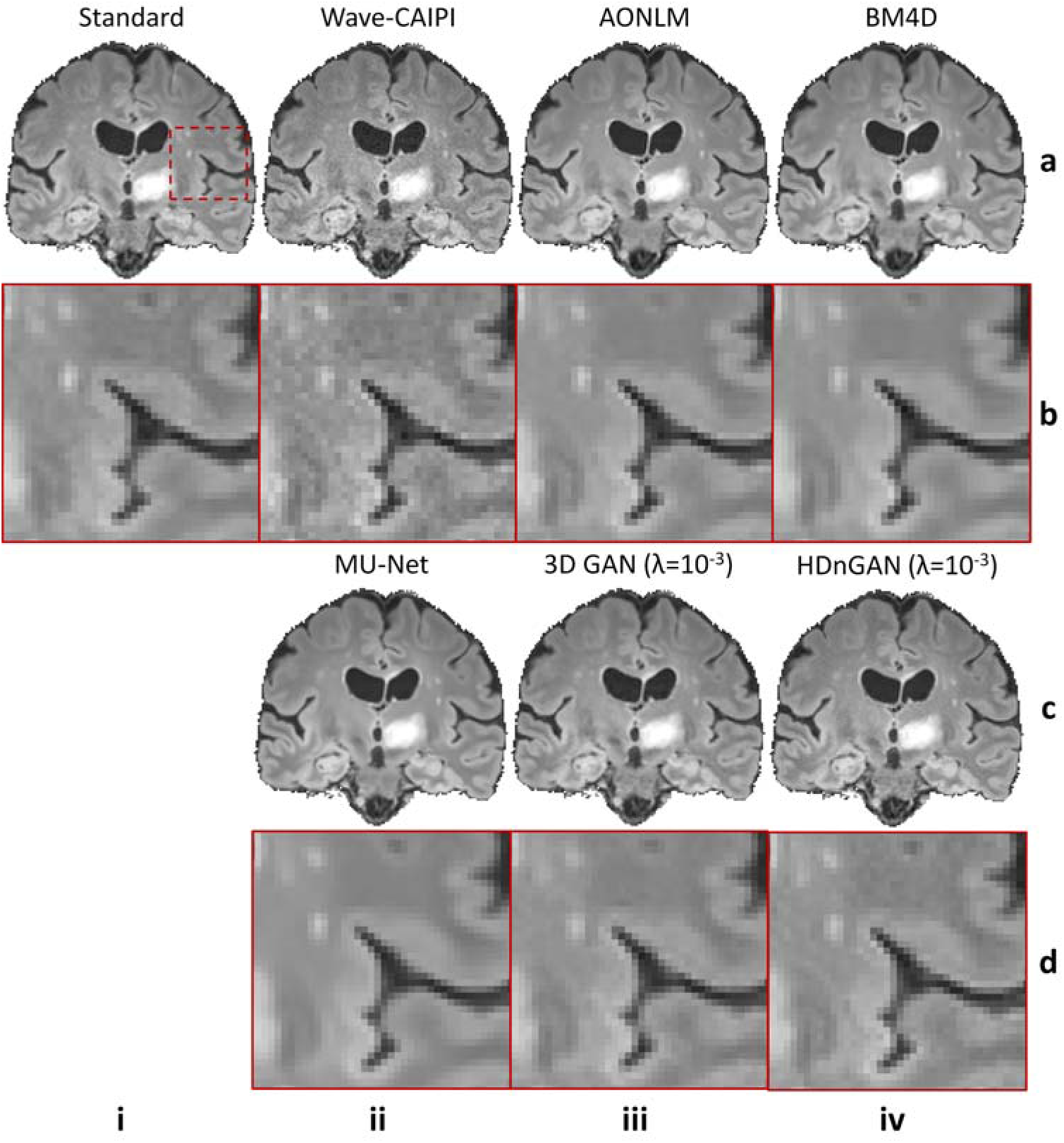
Visual comparison of coronal image slices from different methods. Representative coronal image slices (row a and c) and enlarged regions (row b and d) from standard FLAIR data (row a, b, column i), Wave-CAIPI data (row a, b, column ii), AONLM-denoised results (row a, b, column iii), BM4D-denoised results (row a, b, column iv), MU-Net results (row c, d, column ii), 3D GAN (*λ* = 10^−3^) results (row c, d, column iii) and HDnGAN (*λ* = 10^−3^) results (row c, d, column iv) from one evaluation subject.

**Supplementary Information Figure 5.**
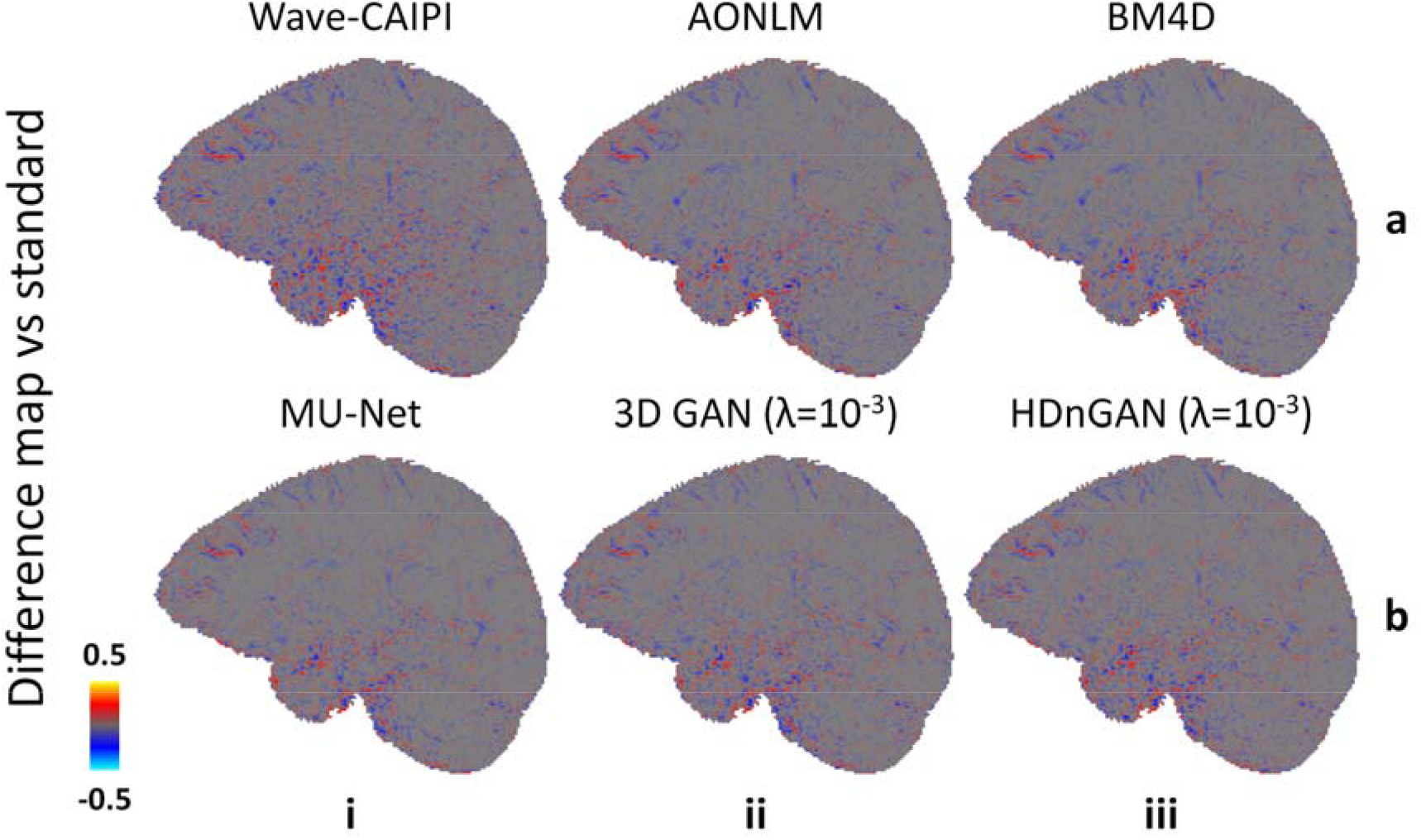
Difference maps. Maps of the difference between images from different methods and standard FLAIR images shown for a representative sagittal image slice from an evaluation subject.

**Supplementary Information Figure 6.**
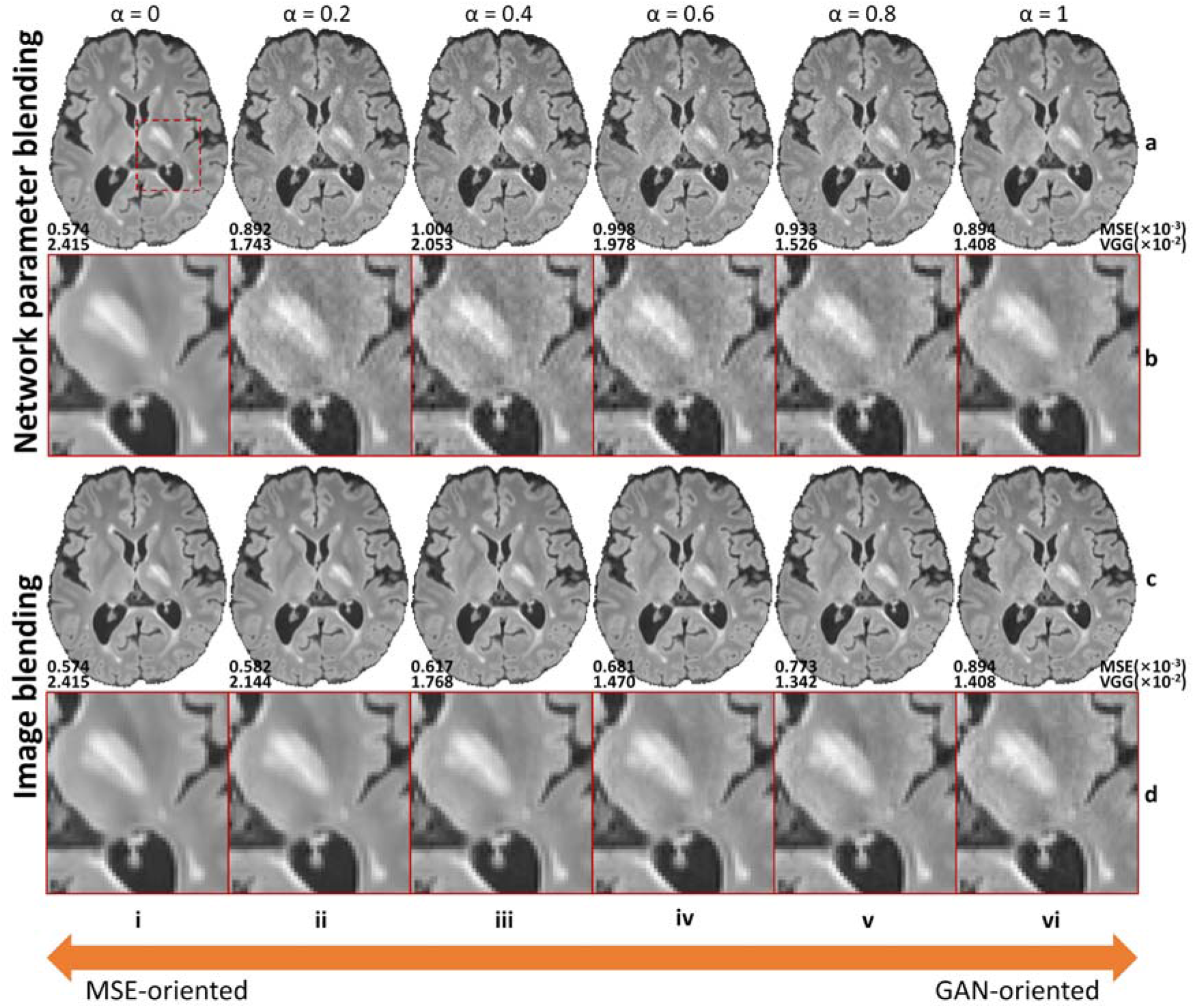
Effects of network parameter blending and image blending on image quality. Representative axial image slices (rows a and c) and enlarged views of the left basal ganglia and left thalamus (rows b and d) from GAN generator network parameter blending (rows a and b) and GAN-synthesized image blending (rows c and d) with different weights (α) from a multiple sclerosis patient for evaluation. For α = 0 (column i), the results are from mean squared error (MSE)-oriented training (i.e., only minimize the mean voxel-wise squared error). In this case, the network is effectively the generator (i.e., MU-Net). For α = 1 (column vi), the results are from GAN-oriented training (i.e., only minimize the adversarial loss). Intermediate results are either synthesized by a generator network whose parameters at each layer are a weighted summation of network parameters from the MSE-orientated training and GAN-oriented training (i.e., network parameter blending, rows a and b), or a voxel-wise weighted summation of synthesized images from the MSE-orientated training and GAN-oriented training (i.e., image blending, rows c and d). Image similarity metrics including the mean squared error (MSE) and VGG perceptual loss (VGG) are listed to quantify the similarity between images from different methods and the standard FLAIR image.

**Supplementary Information Figure 7.**
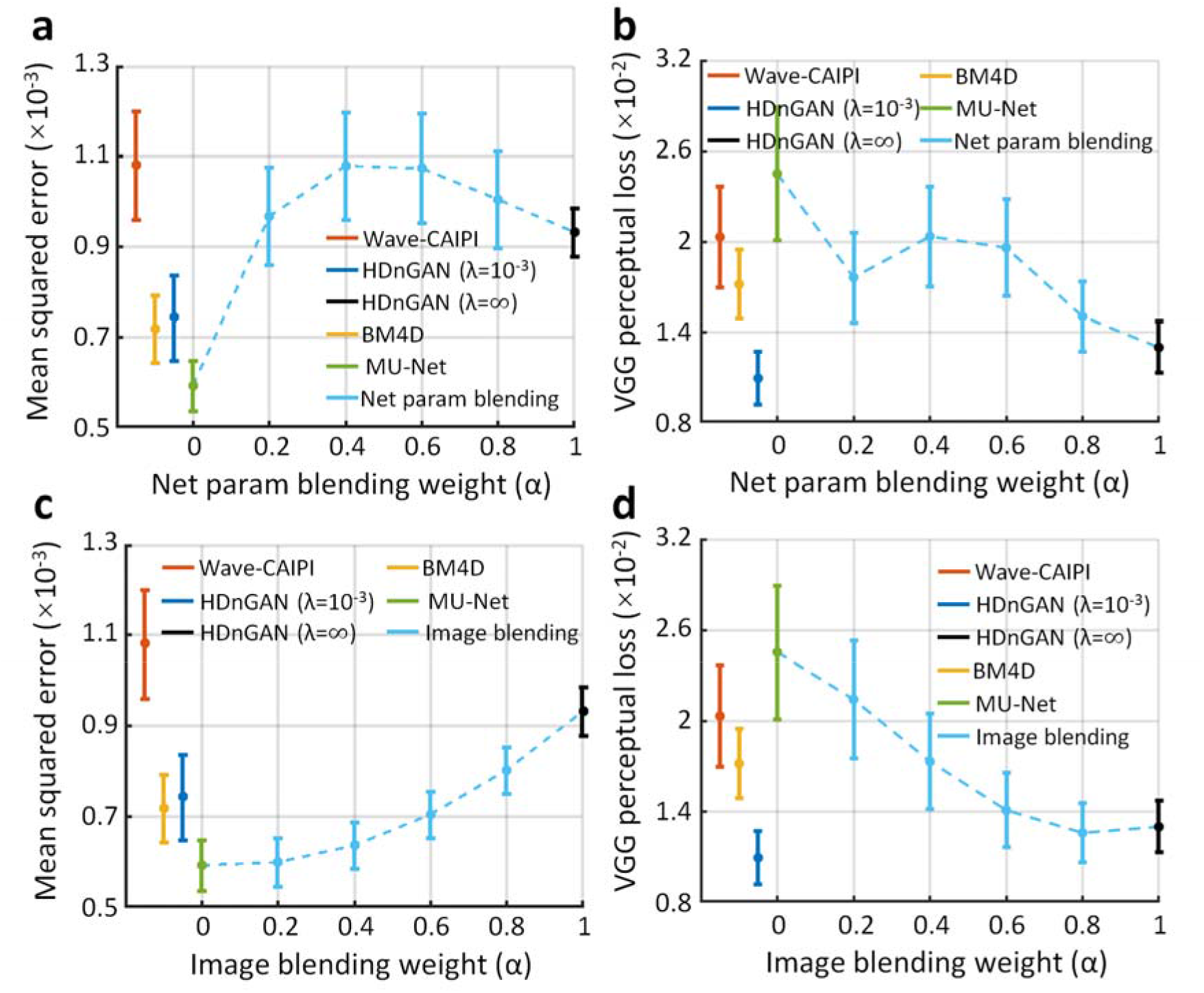
Quantification of the effects of network parameter blending and image blending. The similarity between images derived from different methods and the standard FLAIR images is quantified using the mean squared error (MSE) (a and c) and VGG perceptual loss (b and d). The red, yellow, blue, green, black, and light blue dots and error bars represent the group mean and group standard deviation of different metrics for Wave-CAIPI images, BM4D results, HDnGAN (*λ* = 10^−3^) results, MU-Net results, HDnGAN (*λ* = ∞) results and results of GAN generator parameter blending (a and b) or GAN-synthesized image blending (c and d) with different weights (α). The metrics were calculated from eight patients for evaluation.

**Supplementary Information Table 1.** Comparison of scores from the multi-reader study. Raw *P* values from t-tests assess whether there are significant differences between image quality scores from different methods on the eight evaluation subjects from two neuroradiologists. Raw *P* values surviving multiple comparisons correction at an FDR threshold of 0.05 are marked in bold. All *P* values are two-sided.

|  | Sharpness | SNR | Lesion conspicuity | Overall |
| --- | --- | --- | --- | --- |
| Standard vs Wave-CAIPI | 4.10E-2 | **1.19E-9** | **1.42E-2** | **1.41E-3** |
| Standard vs AONLM | **1.09E-4** | 6.69E-1 | **1.03E-3** | **1.40E-2** |
| Standard vs BM4D | **5.54E-9** | 7.51E-1 | **1.03E-3** | **4.59E-5** |
| Standard vs MU-Net | **2.32E-7** | 7.91E-1 | **4.19E-4** | **1.53E-5** |
| Standard vs 3D GAN ($\lambda=10^{-3}$) | **2.88E-7** | **1.97E-2** | **9.37E-5** | **4.28E-6** |
| Standard vs HDnGAN ($\lambda=10^{-3}$) | 4.10E-2 | 3.33E-1 | 2.75E-2 | 3.33E-1 |
| Wave-CAIPI vs AONLM | 4.51E-1 | **2.88E-7** | 5.12E-1 | 3.88E-1 |
| Wave-CAIPI vs BM4D | **3.45E-3** | **1.66E-4** | 4.26E-1 | 4.85E-1 |
| Wave-CAIPI vs MU-Net | **2.12E-2** | **3.93E-4** | 8.51E-2 | 6.86E-2 |
| Wave-CAIPI vs 3D GAN ($\lambda=10^{-3}$) | **1.41E-3** | **2.54E-5** | 7.15E-2 | 3.33E-1 |
| Wave-CAIPI vs HDnGAN ($\lambda=10^{-3}$) | 3.00E-1 | **1.13E-8** | 6.12E-1 | **2.76E-3** |
| AONLM vs BM4D | 2.95E-2 | 4.32E-1 | 1.00E-0 | 1.40E-2 |
| AONLM vs MU-Net | **1.07E-3** | 1.00E-0 | 7.15E-2 | **6.62E-3** |
| AONLM vs 3D GAN ($\lambda=10^{-3}$) | 8.95E-2 | **8.97E-3** | **4.20E-4** | 2.95E-2 |
| AONLM vs HDnGAN ($\lambda=10^{-3}$) | **1.52E-2** | 6.69E-1 | 1.74E-1 | 9.62E-2 |
| BM4D vs MU-Net | 3.83E-1 | 4.32E-1 | 4.01E-2 | 1.64E-1 |
| BM4D vs 3D GAN ($\lambda=10^{-3}$) | 1.00E-0 | 1.04E-1 | 5.48E-2 | 1.00E-0 |
| BM4D vs HDnGAN ($\lambda=10^{-3}$) | **4.21E-7** | 3.33E-1 | 5.48E-2 | **1.09E-4** |
| MU-Net vs 3D GAN ($\lambda=10^{-3}$) | 4.23E-1 | 5.44E-2 | 3.36E-1 | **4.10E-2** |
| MU-Net vs HDnGAN ($\lambda=10^{-3}$) | **1.53E-5** | 7.74E-1 | **1.05E-3** | **1.36E-5** |
| 3D GAN ($\lambda=10^{-3}$) vs HDnGAN ($\lambda=10^{-3}$) | **9.42E-6** | **3.83E-3** | **7.28E-5** | **7.01E-6** |

